# Sex Chromosome Heteromorphism and the Fast-X Effect in Poeciliids

**DOI:** 10.1101/2021.09.03.458929

**Authors:** Iulia Darolti, Lydia J. M. Fong, Benjamin A. Sandkam, David C. H. Metzger, Judith E. Mank

## Abstract

Accelerated rates of sequence evolution on the X chromosome compared to autosomes, known as Fast-X evolution, have been observed in a range of heteromorphic sex chromosomes. However, it remains unclear how early in the process of sex chromosome differentiation the Fast-X effect becomes detectible. Recently, we uncovered an extreme variation in sex chromosome heteromorphism across Poeciliid fish species. The common guppy, *Poecilia reticulata*, Endler’s guppy, *P. wingei*, swamp guppy, *P. picta*, and para guppy, *P. parae*, appear to share the same XY system and exhibit a remarkable range of heteromorphism. This sex chromosome system is absent in recent outgroups. We combined analyses of sequence divergence and polymorphism data across Poeciliids to investigate X chromosome evolution as a function of hemizygosity and reveal the causes for Fast-X effects. Consistent with the extent of Y degeneration in each species, we detect higher rates of divergence on the X relative to autosomes, a signal of Fast-X evolution, in *P. picta* and *P. parae*, while no change in the rate of evolution of X-linked relative to autosomal genes in *P. reticulata*. In *P. wingei*, the species with intermediate sex chromosome differentiation, we see an increase in the rate of nonsynonymous substitutions on the older stratum of divergence only. We also use our comparative approach to test different models for the origin of the sex chromosomes in this clade. Taken together, our study reveals an important role of hemizygosity in Fast-X and suggests a single, recent origin of the sex chromosome system in this clade.

## Introduction

Owing to their unusual inheritance pattern and hemizygosity in males, X chromosomes display many distinct evolutionary properties compared to the rest of the genome (Charlesworth et al. 1987; Vicoso and Charlesworth 2006). The strength of selection and genetic drift, as well as the role of dominance and effective population size, are expected to differ between sex chromosomes and autosomes (Kirkpatrick and Hall 2004; Mank et al. 2010; Meisel and Connallon 2013). Comparing differences in the evolution of sex-linked and autosomal loci is important for understanding patterns of mutation and selection acting across the genome.

An elevated rate of coding sequence evolution on the X chromosome relative to the autosomes, referred to as the Fast-X effect (or Fast-Z in the case of ZW systems) has been observed in a diversity of organisms, including humans (Lu and Wu 2005), primates (Stevenson et al. 2007), mice (Kousathanas et al. 2014), birds (Mank et al. 2007; Mank et al. 2010; Wright et al. 2015), *Drosophila* (Mank et al. 2010; Ávila et al. 2014; Charlesworth et al. 2018), aphids (Jaquiéry et al. 2018), spiders (Bechsgaard et al. 2019), and lepidoptera (Pinharanda et al., 2019; Sackton et al., 2014). However, the magnitude of this effect can vary substantially across species due to demographic factors, differences in mating system or regulatory mechanisms acting on the sex chromosomes (Mank et al. 2010). Notably, all the cases of Fast-X above were observed in highly heteromorphic sex chromosomes, where very few genes, if any, remain on the Y chromosome and the X is largely hemizygous in the heterogametic sex. As such, it remains unclear how early in the process of sex chromosome differentiation the Fast-X effect becomes detectible.

There are two potential causes of Fast-X. The single functional copy of the X chromosome in males leads to hemizygous exposure of genes and therefore stronger purifying selection against recessive deleterious mutations and positive selection for recessive beneficial ones expressed in males (Charlesworth et al. 1987). This adaptive cause of Fast-X is mainly expected in species with heteromorphic sex chromosomes and highly degenerated Y chromosomes, as a large proportion of X-linked genes will be hemizygous in males. Alternatively, in every male and female pair, there are only three X chromosomes to four copies of each autosome, although this varies substantially based on mating system and type of heterogamety (Vicoso and Charlesworth 2009; Mank et al. 2010; Wright and Mank 2013; Wright et al. 2015). This reduced effective population size of the X relative to the autosomes diminishes the relative power of selection on the X, potentially leading to both greater genetic drift and fixation of weakly deleterious mutations on the X chromosome (Charlesworth et al. 1987). This non-adaptive cause of Fast-X does not necessarily require male hemizygosity, just recombination suppression between the X and Y chromosomes and differences in effective population size between X-linked and autosomal loci, and could apply to X loci where the Y copy is expressed and remains functional.

Recently, we uncovered a case of extreme variation across Poeciliids in the rate of sex chromosome degeneration and dosage compensation. The same chromosome pair acts as the XY system in the common guppy, *Poecilia reticulata*, its sister species Endler’s guppy, *P. wingei*, the more distantly related swamp guppy, *P. picta*, and the para guppy, *P. parae*, all of which last shared a common ancestor roughly 20 million years ago (Darolti et al. 2019; Sandkam et al. 2021). Notably, *P. latipinna* and *G. holbrooki*, recent outgroup species to this clade, do not show evidence of a sex chromosome system on the same chromosome (Darolti et al. 2019), and therefore we hypothesized based on parsimony that the sex chromosome system arose once, roughly 20 mya (supplementary Fig. S1, Parsimony Model). Since this origin, the sex chromosomes of *P. reticulata* and *P. wingei* have remained largely homomorphic, although *P. wingei* shows greater divergence in a limited region (Darolti et al. 2019). In contrast, the *P. picta* and *P. parae* sex chromosomes are completely nonrecombining (with the exception of a small pseudoautosomal region at the distal end, 20-21Mb, which comprises ∼6% of the chromosome) and the Y chromosome has undergone substantial degeneration, leaving most of the X chromosome in males effectively hemizygous (Darolti et al. 2019; Sandkam et al. 2021). Moreover, *P. picta* and *P. parae* exhibit complete dosage compensation (Darolti et al. 2019; Metzger et al. 2021), which is predicted to accentuate the Fast-X effect (Charlesworth et al. 1987; Mank et al. 2010). The range of sex chromosome differentiation present in this clade allows us to investigate patterns of X chromosome coding sequence evolution and to test causes underlying Fast-X effects.

In contrast to the Parsimony Model, it has recently been proposed that the sex chromosomes of *P. reticulata* represent a recent turnover event, and that the highly diverged *P. picta* and *P. parae* system is in fact ancient (Charlesworth et al. 2021). This Turnover Model (supplementary Fig. S1) posits that the *P. picta* X chromosome arose well before the immediate ancestor of *P. parae, P. picta, P. reticulata* and *P. wingei* (Fig. 1 from Charlesworth et al. 2021), and that the almost complete loss of Y gene content and evolution of dosage compensation observed in *P. picta* occurred well before the species split. This model supposes that the homomorphism observed in *P. reticulata* and *P. wingei* represents a turnover event, where the current guppy X and Y chromosomes arose from the ancestral X of *P. picta*. Sex chromosome turnovers are frequent in many animal groups (Bachtrog et al. 2014), so the model is plausible. Disentangling the evolutionary origin of this sex chromosome pair is important for understanding the causes of the observed heterogeneity in sex chromosome differentiation patterns and dosage compensation in this clade.

**Figure 1.**
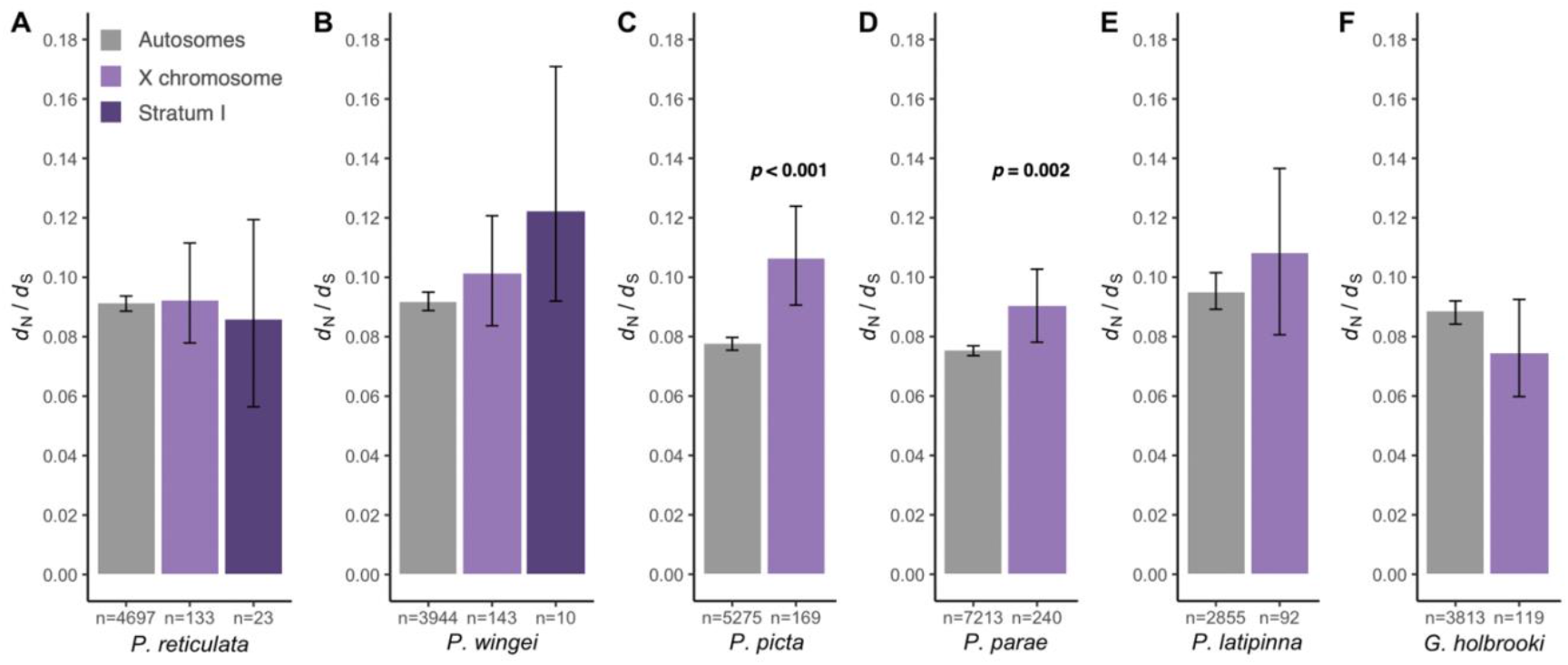
Estimates of the rate of divergence (*d*_N_/*d*_S_) for autosomal and X-linked genes in (A) *P. reticulata*, (B) *P. wingei*, (C) *P. picta*, (D) *P. parae*, (E) *P. latipinna* and (F) *G. holbrooki*. For all species, the X chromosome category excludes genes on the PAR. Additionally, in *P. reticulata* and *P. wingei*, the X chromosome category excludes genes in Stratum I. In *P. latipinna* and *G. holbrooki*, the X chromosome represents the chromosome syntenic to guppy chromosome 12. 95% confidence intervals are based on bootstrapping with 1,000 replicates. Differences between autosomal and X-linked loci are based on 1,000 replicate permutation tests. Only significant differences are shown (*p* value < 0.01).

The unique molecular and evolutionary signatures that accumulate on sex chromosomes can be detected even after they revert to being autosomes. For example, the *Drosophila* dot chromosome was ancestrally a highly differentiated sex chromosome that reverted to an autosome with the emergence of Drosophilidae, and yet it still maintains many of the unique characteristics of a differentiated X chromosome, such as a feminizing effect, non-random gene content and a chromosome-specific gene expression regulatory mechanism (Vicoso and Bachtrog 2013). This means that if the Turnover Model is true, Fast-X patterns, evolving over long periods of time in the distant ancestor of *P. picta* would be evident in internal branches of the phylogeny and should also remain detectible on the X chromosome in *P. reticulata* and *P. wingei* after turnover. Furthermore, the Turnover Model requires additional turnover events in several other taxa within the clade that do not share the same sex chromosome as *P. picta* (supplementary Fig. S1), and we might also expect these species to exhibit ancient patterns of Fast-X on the chromosome, which is fully autosomal. Our comparative dataset allows us to also test these alternative hypotheses for the origin of the guppy sex chromosomes.

Using a combination of sequence divergence, polymorphism and expression data analyses across Poeciliid species, we estimate rates of gene sequence evolution across the genome and assess the presence of Fast-X evolution in each system. Consistent with the extent of Y degeneration, we find significantly higher rates of X coding sequence evolution compared to the autosomes, a signal of Fast-X effect, in *P. picta* and *P. parae*, potentially accelerated by the evolution of chromosome-wide dosage compensation in these species. The absence of a consistent Fast-X evolution in *P. reticulata, P. wingei, P. latipinna*, and *G. holbrooki*, as well as in the ancestral branch to *P. reticulata* and *P. picta*, supports the Parsimony Model.

## Results

### Fast-X evolution in the heteromorphic sex chromosomes of *P. picta* and *P. parae*

We first assessed the strength of Fast-X in our study species with heteromorphic sex chromosomes. The extensive X chromosome hemizygosity of *P. picta* and *P. parae* males is expected to accentuate the strength of Fast-X evolution in these species (Vicoso and Charlesworth 2006), as is the mechanism of complete X chromosome dosage compensation (Mank et al. 2010). Indeed, our analysis in *P. picta* and *P. parae* revealed a higher rate of nonsynonymous substitutions for X-linked loci, excluding genes in the pseudoautosomal region (PAR) (see methods), compared to the autosomes (1,000 replicates permutation test, *p* < 0.001; supplementary Table S1). Both species also exhibit a significantly elevated rate of divergence (*d*_N_/*d*_S_) on the X chromosome relative to the rest of the genome (Fig. 1C, D), consistent with a signal of Fast-X evolution (Fig. 2; supplementary Fig. S2) measured as the ratio of the rate of nonsynonymous over synonymous substitutions for the X chromosome relative to that for the autosomes. Our results thus indicate that the Fast-X signal is present in both *P. picta* and *P. parae*.

**Figure 2.**
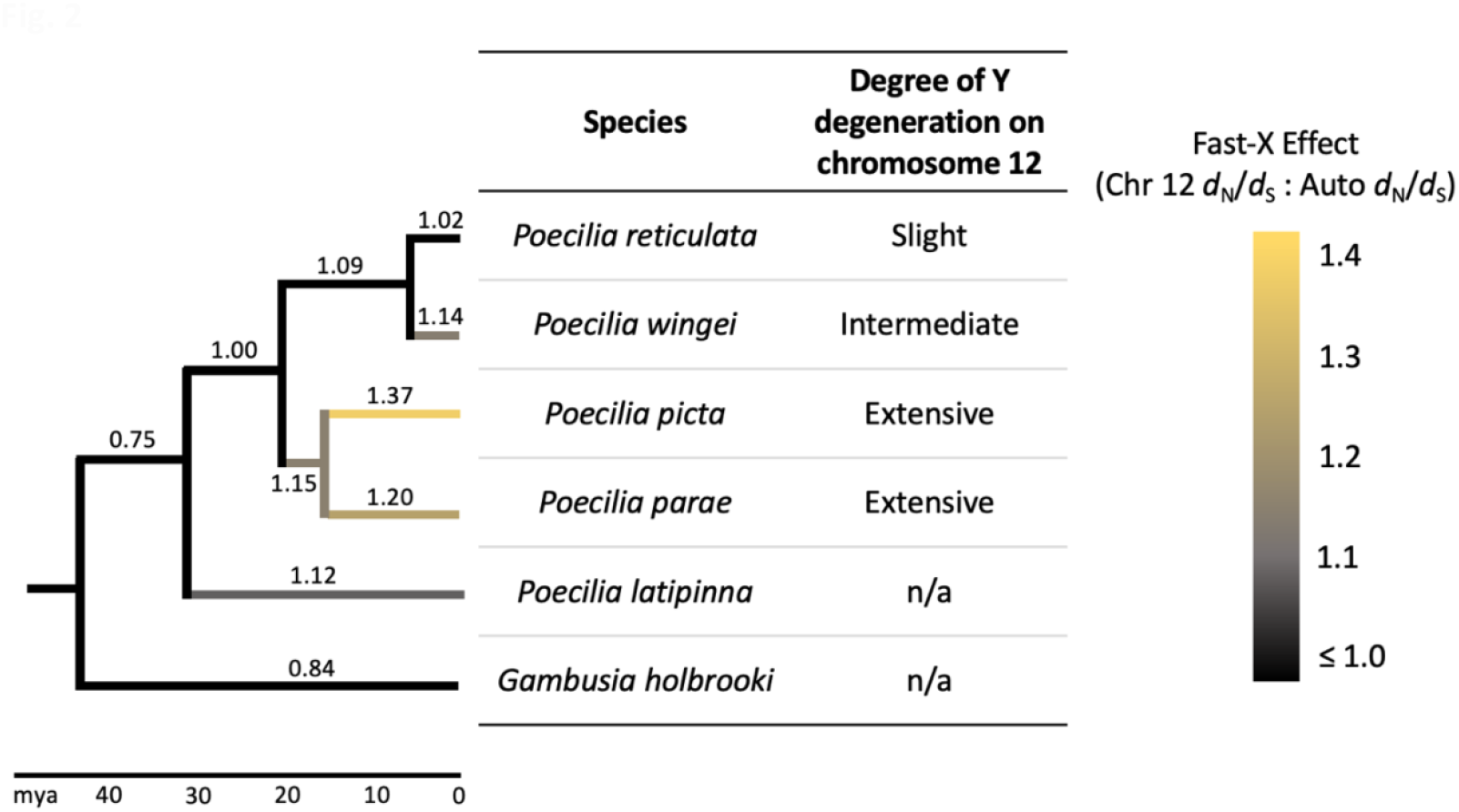
Fast-X effect, calculated as the ratio of *d*_N_/*d*_S_ for the X chromosome (excluding the PAR) to that of the autosomes, across the poeciliids. In *P. latipinna* and *G. holbrooki*, chromosome 12 has not been implicated as the sex chromosome. Numbers on each branch represent the estimated Fast-X effect. Phylogeny based on Meredith et al. 2010 and Rabosky et al. 2018.

### Absence of Fast-X effect in species with homomorphic sex chromosomes

We next asked whether patterns of coding sequence evolution are different between the species with heteromorphic sex chromosomes, *P. picta* and *P. parae*, and the species with homomorphic sex chromosomes, *P. reticulata* and *P. wingei*. Previous work has indicated that *P. reticulata* and *P. wingei* share the same male-heterogametic sex chromosome system (Nanda et al. 2014; Morris et al. 2018; Darolti et al. 2019). Analyses of coverage differences between males and females indicate that Y degeneration is restricted to the distal end of the sex chromosomes, in a region ancestral to *P. reticulata* and *P. wingei* (designated as Stratum I) (Wright et al. 2017; Darolti et al. 2019; Darolti et al. 2020; Fraser et al. 2020; Qiu et al. 2022), and suggest that Y degeneration is slightly more exaggerated in *P. wingei* compared to *P. reticulata*. This nonrecombining region coincides with the previously mapped location of the sex determining region in *P. reticulata* (Winge 1922; Winge 1927; Winge and Ditlevsen 1947; Traut and Winking 2001; Tripathi et al. 2009) and is also found across six natural guppy populations from Trinidad (Almeida et al. 2021).

We therefore assessed the signal of Fast-X evolution for genes in Stratum I (20-26Mb on *P. reticulata* chromosome 12; 17-20Mb on *P. wingei* chromosome syntenic to the guppy sex chromosome). In *P. reticulata*, the rate of divergence for X-linked genes in Stratum I was not significantly different than that for autosomal genes (Fig. 1A; supplementary Table S1). To exclude the possibility that we were lacking power in our analysis due to the small number of genes identified in Stratum I, we reanalyzed *P. reticulata* using Ensembl coding sequences instead of our *de novo* generated transcripts. We were able to recover more than twice as many X-linked loci, however the estimates of divergence remain the same as those based on *de novo* transcripts (supplementary Fig. S3).

In *P. wingei*, genes in Stratum I show significantly higher rates of nonsynonymous and of synonymous substitutions (supplementary Table S1), however the overall rate of divergence is similar between the autosomes and Stratum I (Fig. 1B). *P. wingei* therefore shows an intermediate pattern of Fast-X evolution, not as accentuated as in *P. picta* but more evident than in *P. reticulata* (Fig. 2; supplementary Fig. S2). Recent work has suggested that the *P. wingei* X and Y chromosomes are somewhat more differentiated from each other compared to those of *P. reticulata* (Darolti et al. 2019), and there is also evidence for this from previous cytogenetic work (Nanda et al. 1992; Nanda et al. 2014). This greater divergence might explain the difference in *d*_N_/*d*_S_ estimates for X-linked genes in Stratum I that we observe between these two species, though the effect is negligible. However, it should be noted that following filtering (see methods), only a handful of *P. wingei* genes remained in Stratum I, and as such our statistical power to detect a significant Fast-X signal in this species is reduced.

If the ancestor of the guppy and *P. picta* had an almost completely degenerated sex chromosome system and had evolved dosage compensation (as suggested in the model by Charlesworth et al. 2021), then a signal of Fast-X evolution would have also accumulated prior to the species split. The molecular and evolutionary signatures that accumulate on sex chromosomes remain observable even when they revert to being autosomes (Vicoso and Bachtrog 2013). As such, if the guppy sex chromosomes represent a turnover event, we would expect the pattern of Fast-X that we observed in *P. picta* and *P. parae* to still be observed throughout the *P. reticulata* and *P. wingei* sex chromosomes outside of Stratum I. Thus, we next estimated rates of sequence divergence for the remainder of the X chromosome, excluding genes in Stratum I and genes on the PAR (see methods), as the high recombination events in the PAR can alter rates of evolution and polymorphism (Otto et al. 2011). Although the Fast-X effect was >1 in *P. wingei* (Fig. 2; supplementary Fig. S2), the rate of mutation and mean *d*_N_/*d*_S_ on the X chromosome were not different from those on the autosomes in either *P. reticulata* or *P. wingei* (Fig. 1A, B; supplementary Table S1). This finding is consistent with the limited Y degeneration observed in these species (Wright et al. 2017; Darolti et al. 2019), and does not support the Turnover Model.

Previous comparisons of replicate *P. reticulata* natural population have shown evidence for greater X-Y divergence in replicate upstream, low predation populations relative to their downstream, high predation populations pair within Quare, Aripo and Yarra rivers (Wright et al. 2017; Almeida et al. 2021). This pattern of divergence has occurred over a relatively short time span, at some point after the colonization of Trinidad during the last glacial maximum. Therefore, we investigated whether we can detect higher rates of X chromosome to autosome coding sequence evolution in upstream compared to downstream populations. We found a significantly elevated *d*_N_/*d*_S_ on the X chromosome, excluding genes on the PAR and those on Stratum I, compared to the autosomes in the Yarra upstream population only (supplementary Tables S2-4), and across all rivers we did not find a consistently stronger Fast-X effect in the upstream relative to the downstream populations (Fig. 3). Compared to the other rivers, the upstream population of the Yarra river showed the greatest increase in Y divergence relative to the downstream population, measured by both male:female F_ST_ and male-specific SNPs (Almeida et al. 2021). X-Y divergence between upstream and downstream populations is greater in Yarra compared to Quare and Aripo, and this might explain why a pattern of Fast-X effect in the upstream population was recovered in Yarra alone.

**Figure 3.**
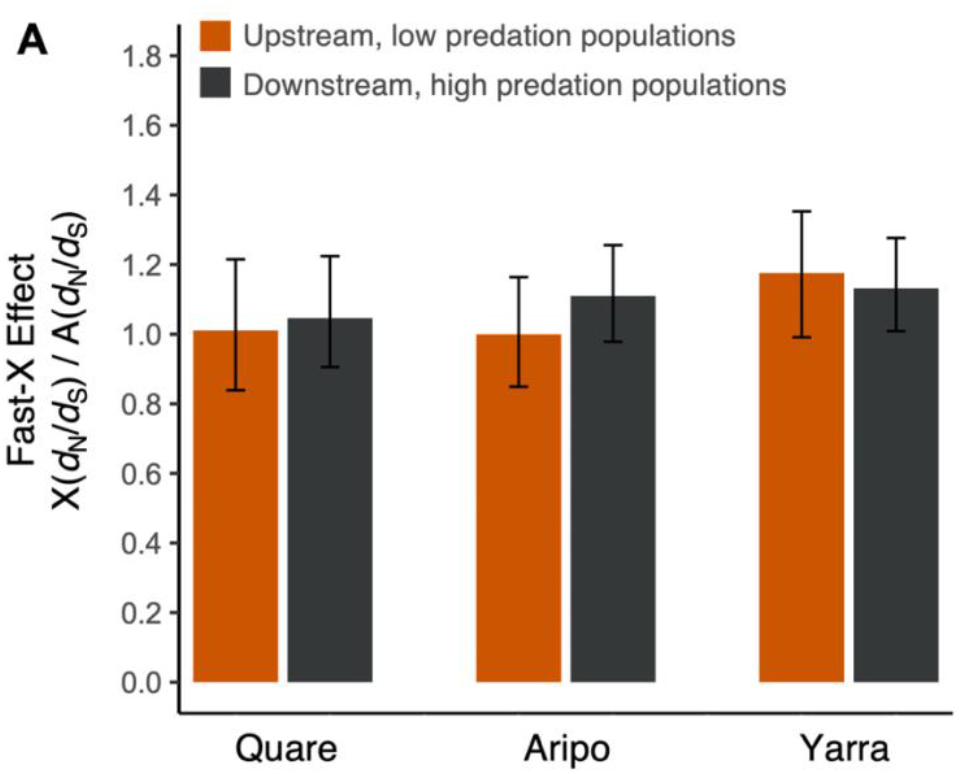
Estimates of rates of evolution in wild *P. reticulata* populations. (A) Fast-X effect across *P. reticulata* upstream, low predation populations (orange) and downstream, high predation populations (black) in the Quare, Aripo and Yarra rivers. The Fast-X effect is calculated as the ratio of the rate of nonsynonymous over synonymous substitutions for the X chromosome, excluding the PAR, relative to that for the autosomes.

### Ancestral Fast-X

If sex chromosome divergence and dosage compensation occurred before the split of *P. picta* from the *P. latipinna* or the *Gambusia* clade, we would expect that an elevated rate of sequence divergence would be seen for X-linked relative to autosomal loci in the branch immediately ancestral to the *P. picta, P. parae, P. reticulata* and *P. wingei* clade. We ran separate phylogenetic analyses with different outgroup species, however neither recovered any difference in the overall rate of divergence or the rates of either nonsynonymous or synonymous substitutions between the autosomal and sex-linked gene categories (Table 1, Fig. 2). This indicates the absence of Fast-X signal in the branch that is ancestral to *P. reticulata, P. wingei* and *P. picta*, which is inconsistent with the Turnover model and the hypothesis of an ancestral degenerated sex chromosome system.

**Table 1.** Divergence estimates for autosomal and X-linked genes on the ancestral branch to *P. reticulata* and *P. picta*. The X chromosome category excludes genes on the PAR. 95% confidence intervals are based on bootstrapping with 1,000 replicates.

| Phylogeny | Category | No.<br>genes | $d_N$<br>(95% CI) | $d_S$<br>(95% CI) | $d_N/d_S$<br>(95% CI) |
| --- | --- | --- | --- | --- | --- |
| (((*P. reticulata*, *P. picta*),<br>*P. latipinna*), *X. maculatus*) | Autosomes | 2,213 | 0.0013<br>(0.0013-0.0014) | 0.0094<br>(0.0092-0.0097) | 0.1422<br>(0.1338-0.1507) |
|  | X chromosome | 82 | 0.0014<br>(0.0010-0.0018) | 0.0096<br>(0.0085-0.0110) | 0.1416<br>(0.1016-0.1937) |
| (((*P. reticulata*, *P. picta*),<br>*G. affinis*), *X. maculatus*) | Autosomes | 3,654 | 0.0049<br>(0.0048-0.0051) | 0.0451<br>(0.0444-0.0457) | 0.1093<br>(0.1055-0.1130) |
|  | X chromosome | 127 | 0.0052<br>(0.0044-0.0061) | 0.0477<br>(0.0441-0.0510) | 0.1096<br>(0.0924-0.1299) |

It has been proposed that the large-scale Y gene loss observed in *P. picta* would have taken considerable amounts of time, much more than the time since *P. picta* split from *P. reticulata* (Charlesworth et al. 2021), estimated at ∼20 mya (Meredith et al. 2010; Rabosky et al. 2018). If that is the case, the Turnover Model also requires additional sex chromosome turnover events in *P. latipinna*, an outgroup species to the *P. reticulata–P. wingei* which has split from *P. parae* only ∼26 mya (Meredith et al. 2010; Rabosky et al. 2018), if not also in the more distantly related *G. holbrooki* (supplementary Fig. S2). We therefore estimated sequence divergence in the outgroups, *P. latipinna* and *G. holbrooki*, both species in which the guppy chromosome 12 has not been implicated as the sex chromosome, and found no evidence of elevated rates of evolution on the chromosome syntenic to the guppy X relative to the rest of the genome in either species (supplementary Table S1). This result, together with our findings in *P. reticulata* and *P. wingei* supports the Parsimony Model.

### Polymorphism and the underlying causes of Fast-X

A higher *d*_N_/*d*_S_ on the X chromosome relative to the rest of the genome could be the result of both increased genetic drift (Charlesworth et al. 1987; Mank et al. 2010) and increased efficacy of selection (Vicoso and Charlesworth 2009). Therefore, we used sequence and polymorphism data together to test for the causes of elevated Fast-X evolution in *P. picta* and *P. parae*. The McDonald-Kreitman test contrasts the number of nonsynonymous and synonymous substitutions with polymorphisms, where an excess of nonsynonymous substitutions relative to polymorphisms is indicative of positive selection (McDonald and Kreitman 1991). Using this test, we detected no X-linked genes with signatures of positive selection in any of the Poeciliid species. However, the McDonald-Kreitman test is very conservative, as it is restricted to genes with sufficient numbers of substitutions and polymorphisms (Begun et al. 2007; Andolfatto 2008), and as such our analysis was limited to a few X-linked contigs.

To increase our statistical power to detect signatures of positive selection, we also used the direction of selection (DoS) test which is less sensitive to low counts (Stoletzki and Eyre-Walker 2011). For each contig, DoS calculates the difference between the proportion of nonsynonymous substitutions and the proportion of nonsynonymous polymorphisms, where a positive DoS indicates adaptive evolution (Stoletzki and Eyre-Walker 2011). Using this approach, we recovered more genes with signatures of positive selection, however the X chromosome was not enriched in these genes compared to the rest of the genome in any of the species (supplementary Table S6).

Since our results did not identify positive selection as the main force underlying the observed Fast-X effect in *P. picta* and *P. parae*, we next used polymorphism data alone to test for a signal of purifying selection acting on the X chromosome. A higher rate of nonsynonymous to synonymous polymorphisms on the X relative to the autosomes is indicative of reduced efficacy of selection to remove mildly deleterious mutations, while a lower rate suggests purifying selection resulting from hemizygous exposure of deleterious mutations in males. Our results show a lower rate of nonsynonymous polymorphisms on the X chromosome in *P. picta* and *P. parae* only (Table 2, 1,000 replicates permutation tests *p* = 0.04 for *P. picta* and *p* = 0.03 for *P. parae*; supplementary Table S5), suggesting that the observed Fast-X effect in these species is influenced by purifying selection acting to remove deleterious mutations on the X (Charlesworth et al. 1987; Vicoso and Charlesworth 2009).

**Table 2.** Polymorphism estimates for autosomal and X-linked genes across Poeciliid species. Only genes with both divergence and polymorphism data were included in this analysis. For all species, the X chromosome category excludes genes on the PAR. 95% confidence intervals are based on bootstrapping with 1,000 replicates. Differences between autosomal and sex-linked categories are based on 1,000 replicate permutation tests and significant differences are indicated by bold values (*p* < 0.01).

| Species | Category | No. genes | $p_N$ (95% CI) | $p_S$ (95% CI) | $p_N/p_S$ (95% CI) |
| --- | --- | --- | --- | --- | --- |
| *P. reticulata* | Autosomes | 2,211 | 0.0012 (0.0011-0.0012) | 0.0110 (0.0105-0.0113) | 0.1054 (0.0980-0.1134) |
|  | X chromosome | 74 | 0.0012 (0.0008-0.0016) | 0.0131 (0.0109-0.0159) | 0.0883 (0.0591-0.1225) |
| *P. wingei* | Autosomes | 934 | 0.0009 (0.0008-0.0010) | 0.0077 (0.0071-0.0082) | 0.1155 (0.1036-0.1275) |
|  | X chromosome | 34 | 0.0008 (0.0004-0.0012) | 0.0103 (0.0080-0.0130) | 0.0791 (0.0411-0.1257) |
| *P. picta* | Autosomes | 1,485 | 0.0006 (0.0006-0.0007) | 0.0051 (0.0048-0.0053) | 0.1191 (0.1082-0.1311) |
|  | X chromosome | 40 | 0.0003 (0.0002-0.0006) | 0.0043 (0.0030-0.0061) | 0.0790 (0.0438-0.1293) |
| *P. parae* | Autosomes | 4,167 | 0.0010 (0.0009-0.0011) | 0.0078 (0.0076-0.0080) | 0.1307 (0.1249-0.1365) |
|  | X chromosome | 123 | 0.0007 (0.0006-0.0009) | **0.0059 (0.0049-0.0068)** | 0.1301 (0.0983-0.1776) |
| *P. latipinna* | Autosomes | 1,041 | 0.0010 (0.0009-0.0011) | 0.0095 (0.0089-0.0100) | 0.1081 (0.1082-0.1311) |
|  | Chromosome syntenic to guppy X | 35 | 0.0016 (0.0010-0.0024) | 0.0111 (0.0081-0.0147) | 0.1452 (0.0832-0.2469) |
| *G. holbrooki* | Autosomes | 1,464 | 0.0014 (0.0013-0.0015) | 0.0155 (0.0146-0.0163) | 0.0872 (0.0800-0.0947) |
|  | Chromosome syntenic to guppy X | 42 | 0.0013 (0.0008-0.0018) | 0.0191 (0.0153-0.0234) | 0.0659 (0.0410-0.0990) |

## Discussion

### The role of hemizygosity in Fast-X

We observe a clear pattern of Fast-X in *P. picta* and *P. parae*, as shown through the significantly higher rate of nonsynonymous substitutions for X-linked loci (supplementary Table S1) and significantly elevated rate of divergence on the X chromosome relative to the rest of the genome (Fig. 1C, D; Fig. 2), consistent with the extensive Y chromosome degeneration in this species. In contrast, we did not observe significant patterns of Fast-X evolution in either *P. reticulata* or *P. wingei*. There is little evidence of loss of Y gene coding sequence in either of these species (Darolti et al. 2019; Darolti et al. 2020), and very few, if any, genes are hemizygously expressed in males, even in the older Stratum I. Interestingly, the *P. wingei* Y is somewhat more diverged than the Y in *P. reticulata* (Nanda et al. 1992; Nanda et al. 2014; Darolti et al. 2019), and we observe a slight, though non-significant Fast-X effect in Stratum I in the former. We also observe an effect in the Yarra upstream population, which shows the greatest X-Y divergence across natural populations in Trinidad (Almeida et al. 2021). All this points to the key role of sex chromosome hemizygosity in driving Fast-X evolution.

Some previous work has failed to detect X-Y divergence based on male:female coverage differences in Stratum I of *Poecilia reticulata* (Bergero et al. 2019; Charlesworth et al. 2020; Kirkpatrick et al. 2022). However, studies that have used similar genomic methods across a range of datasets all recovered patterns consistent with this region (Darolti et al 2019; Wright et al. 2017; Almeida et al. 2020; Fraser et al. 2020; Sigeman et al. 2021), as have studies using male-specific sequence (Morris et al. 2018; Almeida et al. 2020; Darolti et al. 2020). A recent methodological analysis revealed that the detection of Stratum I in *P. reticulata* is largely dependent upon the reference genome used (Darolti et al. 2022), with studies using reference genomes from distant populations or even different species less able to detect subtle patterns of divergence that characterize the Y chromosome in this species. Our detection of a slight, but non-significant pattern of Fast-X in Stratum I of the Yarra *P. reticulata* population as well as in the same region of the *P. wingei* X chromosome gives further weight to the presence of this Stratum.

There are two potential causes of Fast-X evolution. Male heterogamety leads to hemizygous exposure of genes resulting in stronger purifying selection against recessive deleterious mutations and positive selection for recessive beneficial ones expressed in males (Charlesworth et al. 1987). Alternatively, the reduced effective population size of the X relative to the autosomes diminishes the relative power of selection on the X, potentially leading to non-adaptive causes of Fast-X (Charlesworth et al. 1987) which do not necessarily require male hemizygosity, just recombination suppression between the X and Y chromosomes.

In addition to hemizygosity and effective population size, theory predicts that complete dosage compensation, whereby balance in expression between the sex chromosomes and the autosomes is restored, may facilitate a more pronounced Fast-X effect (Charlesworth et al. 1987; Mank et al. 2010). Under this theory, in systems with incomplete dosage compensation, where only a subset of the genes are compensated for but overall expression for X-linked genes is reduced compared to the autosomes in males, X-linked beneficial mutations may have lower expression and weaker phenotypic effects in males, potentially limiting the Fast-X effect (Charlesworth et al. 1987; Mank et al. 2010). *P. picta* and *P. parae* have evolved chromosome-wide dosage compensation, seemingly through a mechanism involving the hyperexpression of the single X in males (Darolti et al. 2019; Metzger et al. 2021), and this may have contributed to the observed signature of Fast-X evolution.

The role of hemizygosity and dosage compensation we observe in *P. picta* and *P. parae* Fast-X suggest an adaptive mechanism, and our polymorphism analysis is consistent with greater purifying selection acting in males to remove recessive deleterious variation. However, although we were unable to differentiate adaptive and non-adaptive causes of Fast-X in our polymorphism data, polymorphism estimates are sensitive to demographic fluctuations (Tajima 1989; Pool and Nielsen 2007) and, thus, it is difficult to determine to what extent the Fast-X pattern in *P. picta* is adaptive using polymorphism data alone. Future work identifying true X-hemizygous loci in males and analyzing their patterns of sequence evolution may prove more revealing.

### Differentiating between models of sex chromosome origin

Two models have been proposed for the origin of the sex chromosome system in this group. The Parsimony Model posits that the sex chromosomes arose once roughly 20 mya (supplementary Fig. S1) in the ancestor of the species that all share it and experienced different rates of Y decay in different sub-clades (Darolti et al. 2019), possibly exacerbated by the evolution of complete dosage compensation in the immediate ancestor of *P. picta* and *P. parae* (Lenormand et al. 2020; Metzger et al. 2021; Sandkam et al. 2021). The Turnover Model proposes that the sex chromosomes of *P. reticulata* represent a recent turnover event, and that the highly diverged *P. picta* and *P. parae* system arose well before the immediate ancestor of *P. parae, P. picta, P. reticulata* and *P. wingei*, and large-scale Y degeneration and complete dosage compensation occurred before the most recent common ancestor of the four species (Charlesworth et al. 2021). Because the molecular signatures that accumulate on sex chromosomes remain observable even when they revert to being autosomes (Vicoso and Bachtrog 2013), we were able to critically test these alternative models with several different approaches.

First, if the guppy sex chromosomes represent a turnover event, we would expect the pattern of Fast-X in *P. picta* and *P. parae* to also be observed throughout the *P. reticulata* and *P. wingei* sex chromosomes outside of Stratum I, as the majority of it would be expected to have accumulated prior to turnover. We failed to recover signatures of Fast-X in *P. reticulata* and *P. wingei* that would be consistent with the Turnover Model, even after investigating multiple datasets (based on *de novo* transcripts of a lab population, replicate wild populations, and Ensembl coding sequences).

Second, the Turnover Model also requires additional sex chromosome turnover events in the outgroup species to the *P. reticulata–P. wingei* clade, and we would expect a Fast-X signal at least in the close outgroup *P. latipinna* if not also in *G. holbrooki*, where the proposed ancient X chromosome is presumably now fully autosomal. We did not observe Fast-X in either of these species.

Finally, if the sex chromosomes were ancient, we would expect a branch-specific signature of Fast-X in the ancestral branch preceding the split of *P. picta, P. parae, P. reticulata* and *P. wingei*. Instead, our results support the Parsimony Model, and suggest that the sex chromosomes originated in the immediate ancestor of the *P. picta–P. parae–P. reticulata–P. wingei* clade and divergence occurred more rapidly in the common ancestor of *P. picta* and *P. parae*, potentially accelerated by the evolution of complete X chromosome dosage compensation in this clade, which reduces selection to maintain Y chromosome expression for dosage sensitive genes (Metzger et al. 2021).

### Concluding Remarks

Taken together, our comparative analyses of divergence and polymorphism data across poeciliids reveal that the degree of Fast-X evolution follows the extent of X hemizygosity in males, emphasizing the important role of male hemizygosity in this evolutionary process. Our results across the broader clade are consistent with a recent origin of the sex chromosome system, as opposed to an ancient origin with turnovers. This indicates that patterns of Fast-X evolution can accumulate rapidly in the evolution of sex chromosomes.

## Materials and Methods

### Sample collection and sequencing

We have previously obtained tissue samples, extracted and sequenced RNA from the tails of three males and three females of *P. reticulata, P. wingei, P. picta, P. parae, P. latipinna* and *G. holbrooki* (BioProject IDs PRJNA353986, PRJNA528814, PRJNA741270) (Wright et al. 2017; Darolti et al. 2019; Metzger et al. 2021). *P. reticulata* samples were obtained from our outbred laboratory population originating from the Quare River in Trinidad (Kotrschal et al. 2013). *P. wingei* samples were collected from our laboratory population established from a strain maintained by a UK fish fancier. *P. picta* and *P. parae* samples were acquired from Guyana and Suriname, while *P. latipinna* and *G. holbrooki* samples were obtained in Florida. All samples were collected in accordance with national and institutional ethical guidelines.

We extracted RNA from each sample using the Qiagen RNeasy Kit, following the instructions of the manufacturer. Library preparation and sequencing were performed at the University of Oxford Wellcome Centre for Human Genetics and Genome Québec, following standard Illumina protocols and using the Illumina HiSeq 4000 and NextSeq platforms. The data were quality assessed using FastQC v0.11.3 (www.bioinformatics.babraham.ac.uk/projects/fastqc) and trimmed with Trimmomatic v0.36 (Bolger 2014), removing adaptor sequences, reads with an average Phred score < 15 in a sliding window of four bases, reads with leading or trailing bases with a Phred score < 3 and reads shorter than 50bp following trimming.

### Identifying orthogroups

For each species, we mapped RNA-seq reads to a previously constructed species-specific female *de novo* genome assembly (Wright et al. 2017; Darolti et al. 2019; Sandkam et al. 2021) using HISAT2 v2.0.4 (Kim et al. 2015), with the exception of *P. latipinna* for which a male genome assembly was used, and a non-redundant set of transcripts in GTF file format was constructed using StringTie v1.2.4 (Pertea et al. 2015). We filtered the transcripts for non-coding RNA (ncRNA) by extracting transcript sequences with BEDtools getfasta (Quinlan and Hall 2010) and removing transcripts with a BLAST hit to ncRNA sequences from *Poecilia formosa* (PoeFor_5.1.2), *Oryzias latipes* (MEDAKA1), *Gasterosteus aculeatus* (BROADS1) and *Danio rerio* (GRCz10) from Ensembl 104 (Flicek et al. 2014). Only genes with positional information on chromosomal fragments were kept for further analyses and the longest isoform for each target gene was selected. Lastly, we applied a minimum expression filter of 2 RPKM in at least half of the samples of each sex (minimum of two out of three individuals of either sex), resulting in 13,306 *P. reticulata*, 15,089 *P. wingei*, 13,156 *P. picta*, 24,446 *P. parae*, 14,468 *P. latipinna* and 21,861 *G. holbrooki* genic sequences.

We obtained coding sequences from the outgroup species *P. formosa* (PoeFor_5.1.2), *Xiphophorus maculatus* (Xipmac4.4.2) and *O. latipes* (MEDAKA1) from Ensembl 104 and extracted the longest isoform for each gene. Separately for each of our target species, we determined orthology across target and outgroup sequences using reciprocal BLASTn v2.7.1 (Altschul et al. 1990) with an e-value cut-off of 10e^-10^ and a minimum percentage identity of 30%. For genes with multiple blast hits, we chose the top hit based on the highest BLAST score. Our analysis resulted in 7,296 *P. reticulata*, 7,253 *P. wingei*, 7,786 *P. picta*, 9,171 *P. parae*, 7,251 *P. latipinna* and 6,477 *G. holbrooki* orthogroups (four-way 1:1 orthologs).

We tested the robustness of the 1:1 ortholog datasets resulting from the reciprocal BLASTn approach by separately inferring orthologous groups using OrthoFinder (Emms and Kelly 2015). More than half of the orthogroups recovered using the OrthoFinder approach are shared with the reciprocal BLASTn approach (supplementary Fig. S4). Restricting the sequence divergence analysis (detailed below) to only orthogroups that are shared between the two approaches revealed similar estimates of rates of divergence to the original estimates based on all orthogroups identified using the reciprocal BLASTn approach (supplementary Fig. S5). We therefore concluded that the reciprocal best-hit datasets were appropriate to use in this case.

### Estimating sequence divergence across orthogroups

For each contig of each orthogroup, we obtained open reading frames using *O. latipes* (MEDAKA1) protein-coding sequences from Ensembl 104 and BLASTx v2.3.0 with an e-value cut-off of 10e^-10^ and a minimum percentage identity of 30%, excluding orthogroups without BLASTx hits or valid protein-coding sequences. We aligned orthologous gene sequences with PRANK v170427 (Löytynoja and Goldman 2008), using the rooted tree (((Target species, *P. formosa), X. maculatus*), *O. latipes*), where the target species was in turn *P. reticulata, P. wingei, P. picta, P. parae, P. latipinna* and *G. holbrooki*. Alignments were then filtered to remove gaps.

To avoid false positive signals of adaptive evolution, poorly aligned or error-rich regions were masked with SWAMP (Harrison et al. 2014). We ran SWAMP twice, first using a threshold of six nonsynonymous substitutions in a window size of 15 codons, and second using a threshold of two and a window size of five. This approach eliminates sequencing errors that cause short stretches of nonsynonymous substitutions as well as alignment errors that cause longer stretches of nonhomologous sequence due to variation in exon splicing or misannotation (Harrison et al. 2014). To select these thresholds, we first applied a range of masking criteria on our datasets. We then ran the branch-site test for positive selection on the target species branches for both the unmasked and masked datasets. We visually inspected the alignments of genes with the highest likelihood ratios and chose the masking criteria that was most efficient at reducing false positive rates. Finally, we discarded orthologs for which the alignment length was shorter than 300 bp following gap removal and masking, as these likely represent incomplete sequences.

To obtain divergence estimates for each orthogroup and calculate mean *d*_N_/*d*_S_ across the target species branch, we used branch model (model=2, nssites=0) in the CODEML package in PAML v4.8 (Yang 2007), using the phylogeny ((Target species#1, *P. formosa*), *X. maculatus, O. latipes*), where the target species was successively *P. reticulata, P. wingei, P. picta, P. parae, P. latipinna* and *G. holbrooki*. To avoid inaccurate divergence estimates dues to mutational saturation and double hits, orthologous genes with *d*_S_ > 2 were removed from subsequent analyses (Axelsson et al. 2008).

For each of the target species, we divided orthologs into autosomal and sex-linked categories based on their chromosomal location. The sex-linked category excluded genes in the previously identified pseudoautosomal regions (0-5Mb and >26Mb of *P. reticulata* chromosome 12; >20Mb of *P. wingei, P. picta, P. parae, P. latipinna* and *G. holbrooki* chromosomes that are syntenic to the guppy chromosome 12 (Darolti et al. 2019; Sandkam et al. 2021)). In addition, it has recently been discovered that the *P. reticulata* reference genome, which served as reference for scaffold ordering and orientation for our *P. reticulata* de novo genome assembly, has a large inversion on the X chromosome that is specific to the guppy strain on which the reference genome assembly was built and is not present in any of our lab or wild-caught samples (Darolti et al. 2020; Almeida et al. 2021). We thus corrected for this inversion when excluding genes from the PAR and assigning genes to the sex-linked category in *P. reticulata*.

Separately for each genomic category, we extracted the number of nonsynonymous substitutions (*D*_N_), the number of nonsynonymous sites (N), the number of synonymous substitutions (*D*_S_) and the number of synonymous sites (S). Taking into account alignment length, we calculated mean *d*_N_ and mean *d*_S_ as the ratio of the number of substitutions across all orthologs in that group divided by the number of sites (*d*_N_ = *D*_N_ /N; *d*_S_ = *D*_S_ /S), thus avoiding bias from short sequences and the issue of infinitely high *d*_N_/*d*_S_ estimates due to very low *d*_S_ (Mank et al. 2007). We identified significant differences in *d*_N_, *d*_S_ and *d*_N_/*d*_S_ between genomic categories by subsampling without replacement the autosomal dataset at the size of the smaller dataset (e.g., X chromosome dataset; Stratum I dataset) using 1,000 replicates, and we used bootstrapping with 1,000 replicates to determine 95% confidence intervals for each divergence estimate. The strength of the Faster-X effect was calculated for each target species as the ratio of the rate of divergence for the X chromosome over the rate of divergence for the autosomes X(*d*_N_/*d*_S_):A(*d*_N_/*d*_S_).

### Additional sequence divergence analyses for *P. reticulata*

For comparison to our estimates of rates of divergence based on *de novo* transcripts, we also estimated rates of sequence evolution in *P. reticulata* using publicly available coding sequences from Ensembl 104 (Guppy_female_1.0_MT). For this analysis, we followed the same steps outlined above in terms of extracting the longest isoform for each gene, identifying orthogroups, aligning target and outgroup sequences, masking and obtaining divergence estimates.

In addition, we have previously constructed high quality *P. reticulata* female *de novo* genome assemblies for replicate upstream and downstream populations of three rivers in Trinidad, Quare, Aripo, and Yarra (Almeida et al. 2021). To each genome assembly, we mapped *P. reticulata* RNA-seq reads from our laboratory population and generated a non-redundant set of transcripts for each wild population following the methods detailed above. For each upstream and downstream population, we then estimated rates of coding sequence evolution for the X chromosome, excluding the PAR, and the autosomes by following the same pipeline described in the previous section.

### Estimating rates of divergence for the ancestral branch to *P. reticulata* and *P. picta*

We performed additional sequence divergence analyses following the same methodology described above but using different phylogenies that allowed us to estimate rates of evolution on the internal branch that is ancestral to *P. reticulata* and *P. picta*. In addition to the *X. maculatus* coding sequences, we obtained sequences for *G. affinis* (ASM309773v1) and *P. latipinna* (P_latipinna-1.0) from Ensembl 105. We used the phylogenies ((*P. reticulata, P. picta*)#1, *P. latipinna, X. maculatus*) and ((*P. reticulata, P. picta*)#1, *G. affinis, X. maculatus*) to obtain divergence estimates in PAML. Similarly, we estimated rates of evolution for all the other internal branches of the tree (Fig. 2).

### Polymorphism data

To obtain polymorphism data, for each target species we first mapped female RNA-seq reads to the genome assembly using the STAR v2.4.2a aligner in two-pass mode (Dobin et al. 2013), and then called SNPs using SAMtools v1.3.1 mpileup (Li et al. 2009) with a minimum base quality of 20 and VarScan v2.3.9 mpileup2snp (Koboldt et al. 2012) with a minimum coverage of two, a minimum average quality of 20, a minimum variant allele frequency of 0.1, *p* value of 0.05 and strand filter set to “on”. We next imposed that the polymorphism dataset also passes the filtering criteria used for calculating rates of sequence divergence. As such, we identified codons for which all sites pass the minimum coverage threshold of 20 in at least half of the individuals (in this case, a minimum of two out of three individuals), there are no alignment gaps following alignment with PRANK, and no ambiguity data (Ns) following masking with SWAMP. To make our dataset compatible for the McDonald-Kreitman test of selection, we only kept genes with both divergence and polymorphism data. We have therefore used the same number of nonsynonymous and synonymous sites in our divergence and polymorphism calculations. We identified whether SNPs were synonymous or nonsynonymous by matching them to the reading frame using custom scripts (available at https://github.com/manklab/Darolti_etal_2022_FastX_evolution). For each genomic category, we calculated mean *p*_N_ and mean *p*_S_ as the ratio of the number of polymorphisms across all orthologs in that group divided by the number of sites (*p*_N_=*P*_N_/N; *p*_S_=*P*_S_/S).

### Testing for selection using divergence and polymorphism data

For each species, we estimated the number of genes evolving under neutral and adaptive evolution using the McDonald-Kreitman test (McDonald and Kreitman 1991). Neutral theory predicts that the ratio of nonsynonymous to synonymous changes within species (*p*_N_/*p*_S_) should be equal to that between species (*d*_N_/*d*_S_) (McDonald and Kreitman 1991). As such, the McDonald-Kreitman test identifies signatures of positive selection, where there is an excess of nonsynonymous substitutions relative to polymorphisms (*d*_N_/*d*_S_>*p*_N_/*p*_S_), and of relaxed purifying selection, where there is a deficit of nonsynonymous substitutions compared to polymorphisms (*d*_N_/*d*_S_<*p*_N_/*p*_S_). To test for deviations from neutrality, for each contig, we used a 2 × 2 contingency table and a Fisher’s Exact Test in R v3.6.2 (R Core Team 2015). The power of the McDonald-Kreitman test is limited with low table counts, therefore we restricted this analysis to genes for which the sum of each row and column in the contingency table was equal to or greater than six (Begun et al. 2007; Andolfatto 2008).

As the McDonald-Kreitman test is a very conservative test, we also used the divergence and polymorphism data to calculate the direction of selection statistic (DoS) (Stoletzki and Eyre-Walker 2011). For each gene, we calculated the difference between the proportion of nonsynonymous substitutions and polymorphisms (DoS=*D*_N_(*D*_N_+*D*_S_)–*P*_N_(*P*_N_+*P*_S_)), where positive DoS values are indicative of positive selection. We used Fisher’s Exact test in R to test for significant differences in the proportion of genes under positive selection between the autosomes and the X chromosome in each species.

Lastly, we used the polymorphism data alone to test for an excess or deficit of nonsynonymous polymorphisms on the X chromosome compared to the autosomes, which would suggest a relaxed constraint or purifying selection, respectively. For this, we concatenated *P*_N_ and *P*_S_ estimates within each species and used Fisher’s Exact test in R to test for significant differences in *P*_N_/*P*_S_ between the autosomes and the X chromosome.

## Acknowledgements

This work was supported by the European Research Council (grant agreement 680951), a Canada 150 Research Chair, and an NSERC Discovery Grant to J.E.M. We thank C. Lacy for the fish illustrations and members of the Mank lab for helpful discussions and suggestions.

## Author Contributions

J.E.M. and I.D. designed the research; I.D., L.F., B.A.S., D.C.H.M. and J.E.M. performed the data analysis and contributed to the writing of the manuscript.

## Data availability

Scripts used for data processing and analysis are available at https://github.com/manklab/Darolti_etal_2022_FastX_evolution.

