## Supplementary Information for "Sex Chromosome Heteromorphism and the Fast-X Effect in Poeciliids"

### Supplementary Results

**Table S1. Divergence estimates for autosomal and X-linked genes across Poeciliid species.** For all species, the X chromosome category excludes genes on the PAR. Additionally, in *P. reticulata* and *P. wingei*, the X chromosome category excludes genes in Stratum I. 95% confidence intervals are based on bootstrapping with 1,000 replicates. Differences between autosomal and sex-linked categories are based on 1,000 replicates permutation tests and significant differences are indicated by bold values ($p < 0.01$).

| Species | Category | No. genes | $d_N$ (95% CI) | $d_S$ (95% CI) | $d_N/d_S$ (95% CI) |
|---|---|---|---|---|---|
| *P. reticulata* | Autosomes | 4,697 | 0.0031 (0.0030-0.0032) | 0.0337 (0.0333-0.0342) | 0.0911 (0.0886-0.0937) |
| | X chromosome | 133 | 0.0035 (0.0030-0.0041) | **0.0379 (0.0350-0.0407)** | 0.0922 (0.0776-0.1115) |
| | Stratum I | 23 | 0.0045 (0.0029-0.0059) | **0.0523 (0.0404-0.0629)** | 0.0857 (0.0564-0.1194) |
| *P. wingei* | Autosomes | 3,944 | 0.0031 (0.0030-0.0032) | 0.0333 (0.0328-0.0339) | 0.0919 (0.0888-0.0950) |
| | X chromosome | 143 | 0.0037 (0.0031-0.0043) | 0.0360 (0.0332-0.0389) | 0.1014 (0.0837-0.1207) |
| | Stratum I | 10 | **0.0074 (0.0051-0.0098)** | **0.0598 (0.0423-0.0816)** | 0.1220 (0.0920-0.1709) |
| *P. picta* | Autosomes | 5,275 | 0.0037 (0.0036-0.0038) | 0.0474 (0.0468-0.0480) | 0.0776 (0.0754-0.0797) |
| | X chromosome | 169 | **0.0053 (0.0046-0.0061)** | 0.0498 (0.0463-0.0539) | **0.1061 (0.0906-0.1239)** |
| *P. parae* | Autosomes | 7,213 | 0.0045 (0.0044-0.0046) | 0.0601 (0.0592-0.0610) | 0.0752 (0.0736-0.0769) |
| | X chromosome | 240 | **0.0053 (0.0046-0.0059)** | 0.0584 (0.0549-0.0624) | **0.0901 (0.0781-0.1027)** |
| *P. latipinna* | Autosomes | 2,855 | 0.0010 (0.0009-0.0010) | 0.0100 (0.0098-0.0103) | 0.0949 (0.0892-0.1015) |
| | Chromosome syntenic to guppy X | 92 | 0.0011 (0.0009-0.0015) | 0.0105 (0.0091-0.0122) | 0.1079 (0.0806-0.1366) |
| *G. holbrooki* | Autosomes | 3,813 | 0.0040 (0.0038-0.0041) | 0.0449 (0.0439-0.0468) | 0.0883 (0.0842-0.0920) |
| | Chromosome syntenic to guppy X | 119 | 0.0032 (0.0026-0.0039) | 0.0433 (0.0399-0.0471) | 0.0744 (0.0598-0.0925) |

**Table S2. Divergence estimates for autosomal and X-linked genes between wild *P. reticulata* Quare populations.** For both populations, the X chromosome category excludes genes on the PAR and genes in Stratum I. 95% confidence intervals are based on bootstrapping with 1,000 replicates. Differences between autosomal and sex-linked categories are based on 1,000 replicates permutation tests and significant differences are indicated by bold values ($p < 0.01$).

| Population | Category | No. genes | $d_N$ (95% CI) | $d_S$ (95% CI) | $d_N/d_S$ (95% CI) |
|---|---|---|---|---|---|
| Upstream, low predation | Autosomes | 3,887 | 0.0029 (0.0028-0.0030) | 0.0333 (0.0328-0.0337) | 0.0866 (0.0840-0.0895) |
| | X chromosome | 112 | 0.0034 (0.0028-0.0040) | 0.0368 (0.0336-0.0401) | 0.0916 (0.0734-0.1122) |
| | Stratum I | 31 | 0.0032 (0.0024-0.0043) | **0.0433 (0.0367-0.0505)** | 0.0748 (0.0564-0.0966) |
| Downstream, high predation | Autosomes | 5,159 | 0.0030 (0.0029-0.0031) | 0.0336 (0.0331-0.0340) | 0.0888 (0.0861-0.0915) |
| | X chromosome | 138 | 0.0032 (0.0028-0.0036) | 0.0343 (0.0317-0.0374) | 0.0936 (0.0790-0.1103) |
| | Stratum I | 33 | 0.0038 (0.0027- 0.0053) | **0.0446 (0.0369-0.0528)** | 0.0865 (0.0662-0.1115) |

**Table S3. Divergence estimates for autosomal and X-linked genes between wild *P. reticulata* Aripo populations.** For both populations, the X chromosome category excludes genes on the PAR and genes in Stratum I. 95% confidence intervals are based on bootstrapping with 1,000 replicates. Differences between autosomal and sex-linked categories are based on 1,000 replicates permutation tests and significant differences are indicated by bold values ($p < 0.01$).

| Population | Category | No. genes | $d_N$ (95% CI) | $d_S$ (95% CI) | $d_N/d_S$ (95% CI) |
|---|---|---|---|---|---|
| Upstream, low predation | Autosomes | 3,890 | 0.0028 (0.0028-0.0029) | 0.0332 (0.0327-0.0337) | 0.0854 (0.0827-0.0880) |
| | X chromosome | 116 | 0.0032 (0.0026-0.0038) | **0.0376 (0.0345-0.0406)** | 0.0850 (0.0683-0.1040) |
| | Stratum I | 37 | 0.0035 (0.0027-0.0045) | **0.0414 (0.0356-0.0481)** | 0.0851 (0.0652- 0.1087) |
| Downstream, high predation | Autosomes | 6,498 | 0.0029 (0.0028-0.0030) | 0.0334 (0.0331-0.0338) | 0.0868 (0.0847-0.0892) |
| | X chromosome | 202 | 0.0033 (0.0029-0.0037) | 0.0336 (0.0315-0.0357) | 0.0984 (0.0844-0.1119) |
| | Stratum I | 39 | 0.0039 (0.0030-0.0050) | **0.0491 (0.0433-0.0559)** | 0.0799 (0.0632-0.0983) |

17

**Table S4. Divergence estimates for autosomal and X-linked genes between wild *P. reticulata* Yarra populations.** For both populations, the X chromosome category excludes genes on the PAR and genes in Stratum I. 95% confidence intervals are based on bootstrapping with 1,000 replicates. Differences between autosomal and sex-linked categories are based on 1,000 replicates permutation tests and significant differences are indicated by bold values ($p < 0.01$).

| Population | Category | No. genes | $d_N$ (95% CI) | $d_S$ (95% CI) | $d_N/d_S$ (95% CI) |
|---|---|---|---|---|---|
| Upstream, low predation | Autosomes | 5,450 | 0.0030 (0.0029-0.0030) | 0.0334 (0.0334-0.0335) | 0.0868 (0.0844-0.0892) |
| | X chromosome | 130 | 0.0034 (0.0028-0.0041) | 0.0308 (0.0286-0.0333) | **0.1100 (0.0905-0.1321)** |
| | Stratum I | 52 | **0.0041 (0.0031-0.0053)** | **0.0525 (0.0466-0.0590)** | 0.0781 (0.0607- 0.0982) |
| Downstream, high predation | Autosomes | 7,822 | 0.0029 (0.0028-0.0029) | 0.0335 (0.0331-0.0338) | 0.0860 (0.0841-0.0880) |
| | X chromosome | 208 | 0.0034 (0.0030-0.0038) | 0.0331 (0.0314-0.0351) | 0.1019 (0.0886-0.1152) |
| | Stratum I | 60 | 0.0036 (0.0028-0.0044) | **0.0459 (0.0413-0.0514)** | 0.0775 (0.0614-0.0943) |

**Table S5. Differences between nonsynonymous and synonymous polymorphisms on the autosomes and X chromosome of each species.** For all species, the X chromosome category excludes genes on the PAR.

| **Species** | **Category** | $P_N$ | $P_S$ | ***p* value**[a] |
|---|---|---|---|---|
| *P. reticulata* | Autosomes | 1,890 | 5,566 | $p = 0.76$ |
| | X chromosome | 59 | 184 | |
| *P. wingei* | Autosomes | 650 | 1,783 | $p = 0.69$ |
| | X chromosome | 18 | 56 | |
| *P. picta* | Autosomes | 837 | 2,178 | $p = 0.24$ |
| | X chromosome | 12 | 47 | |
| *P. parae* | Autosomes | 3,673 | 8,645 | $p = 0.1$ |
| | X chromosome | 83 | 197 | |
| *P. latipinna* | Autosomes | 1,165 | 3,265 | $p = 0.24$ |
| | Chromosome syntenic to guppy X | 52 | 107 | |
| *G. holbrooki* | Autosomes | 1,212 | 4270 | $p = 0.08$ |
| | Chromosome syntenic to guppy X | 28 | 144 | |

[a]*p* values from Fisher's Exact test

**Table S6. Direction of Selection test of positive selection**

| Species | Group | Total no. of genes[a] | No. of genes under positive selection (*p* value[b]) |
|---|---|---|---|
| *P. reticulata* | Autosomes | 2,211 | 928 |
| | X chromosome | 74 | 35 (*p* = 0.60) |
| *P. wingei* | Autosomes | 934 | 402 |
| | X chromosome | 34 | 16 (*p* = 0.76) |
| *P. picta* | Autosomes | 1,485 | 728 |
| | X chromosome | 40 | 25 (*p* = 0.35) |
| *P. parae* | Autosomes | 4167 | 1837 |
| | X chromosome | 123 | 61 (*p* = 0.46) |
| *P. latipinna* | Autosomes | 1,041 | 287 |
| | Chromosome syntenic to guppy X | 35 | 7 (*p* = 0.57) |
| *G. holbrooki* | Autosomes | 1,464 | 658 |
| | Chromosome syntenic to guppy X | 42 | 21 (*p* = 0.68) |

[a]Includes only genes with both divergence and polymorphism data.
[b]*p* values from Fisher's Exact test comparing the proportion of genes with a signature of positive selection between the autosomes and the X chromosome.

| Species | Sex chromosome on guppy chromosome 12 | Degree of Y degeneration on guppy chromosome 12 | Refs |
|---|---|---|---|
| *Poecilia reticulata* | Yes | Slight | 1, 2, 3, 4, 5 |
| *Poecilia wingei* | Yes | Intermediate | 4, 5 |
| *Poecilia picta* | Yes | Extensive | 4 |
| *Poecilia parae* | Yes | Extensive | 6,7 |
| *Poecilia latipinna* | No | n/a | 4 |
| *Gambusia holbrooki* | No | n/a | 4 |

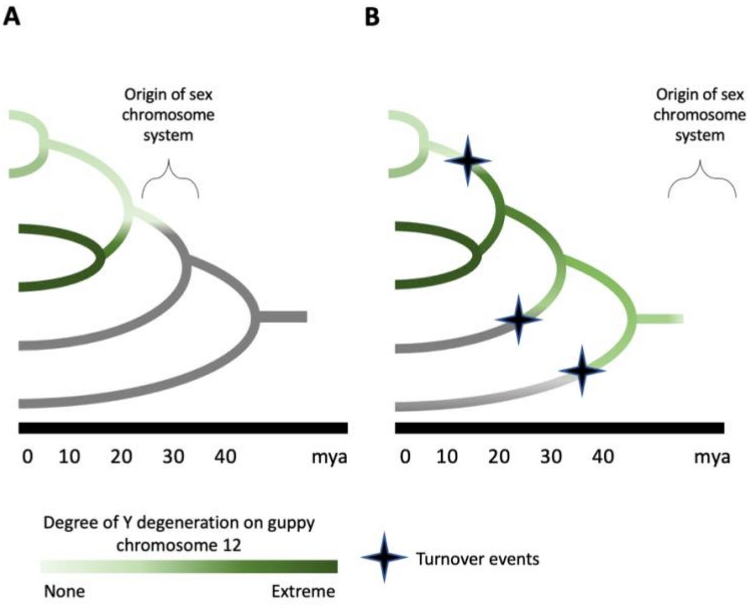


**Fig. S1. Alternative models for the origin of the guppy sex chromosome system.** The Parsimony Model (A) posits that the sex chromosomes originated in the ancestor of *P. reticulata*, *P. wingei*, *P. picta* and *P. parae*, all of which exhibit X-Y divergence on guppy chromosome 12. Once originated, the rate of Y divergence varies across clades and species (shaded in green). The Turnover Model[8] posits that that the sex chromosome system originated much earlier, although the exact date is unspecified, and then exhibited a gradual rate of Y degeneration. This requires turnover events in the ancestor of the *P. reticulata-P. wingei clade*, as well as at least two additional turnover events in the ancestor of *P. latipinna* and *Gambusia holbrooki* (lineages where guppy chromosome 12 has not been implicated as the sex chromosomes are in grey). At this time, no outgroup to the *P. picta-P. parae* group is known to exhibit heteromorphic sex chromosomes on guppy chromosome 12. Phylogeny based on refs 9, 10.

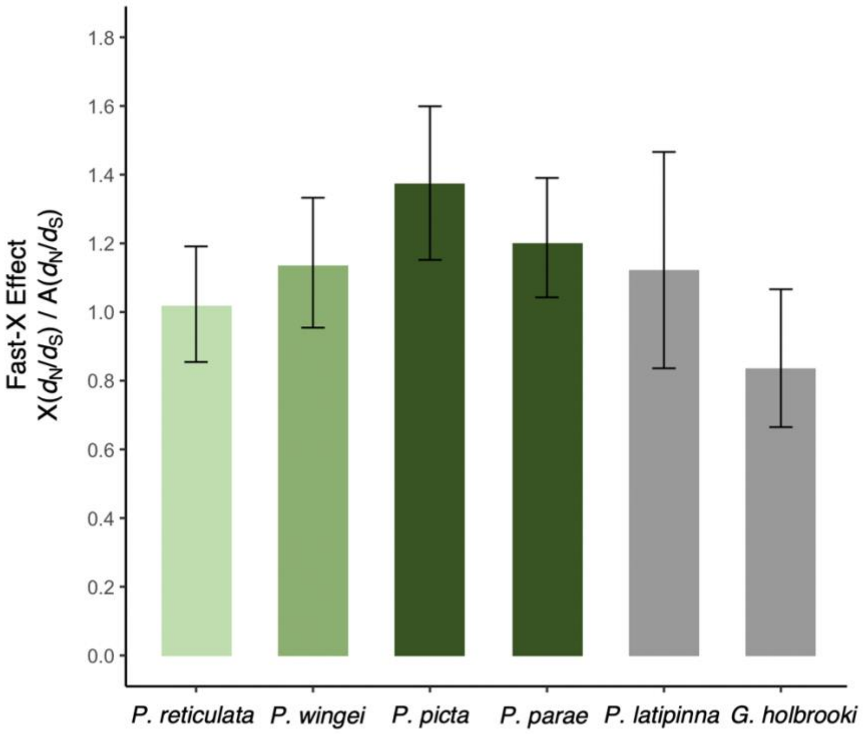


**Fig. S2. Fast-X effect, calculated as the ratio of $d_N/d_S$ for the X chromosome (excluding the PAR) to that of the autosomes, across the poeciliids.** In *P. latipinna* and *G. holbrooki*, the X chromosome represents the chromosome syntenic to guppy chromosome X**.** 95% confidence intervals are based on bootstrapping with 1,000 replicates. Green color shading represents the extent of sex chromosome degeneration (mild degeneration in light green to extreme degeneration in dark green), while species in which the guppy chromosome 12 has not been implicated as the sex chromosome are shown in grey.

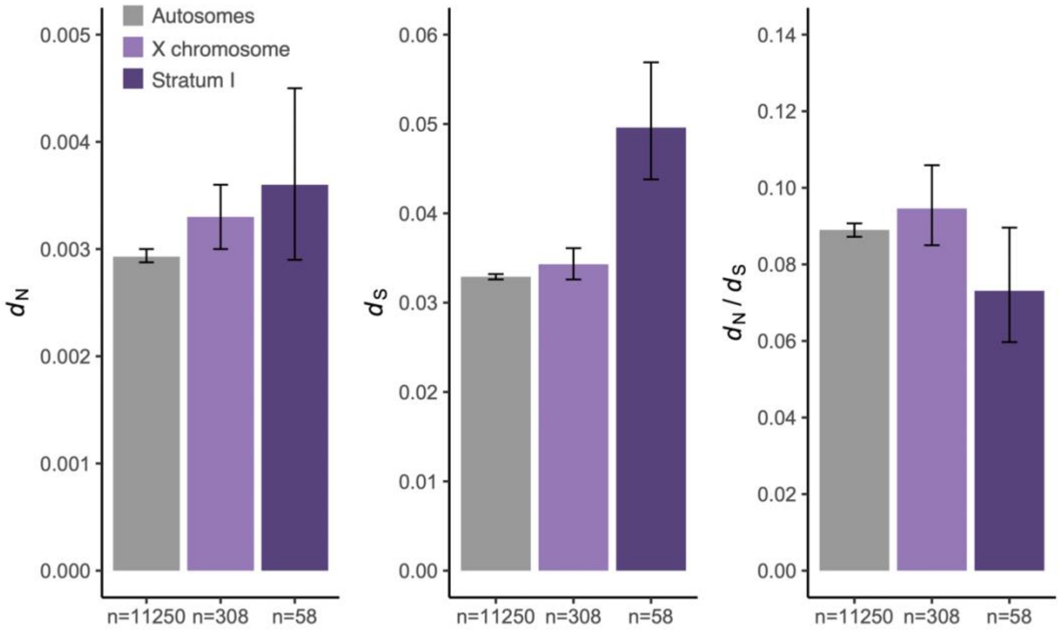


**Fig. S3. Estimates of nonsynonymous substitutions ($d_N$), synonymous substitutions ($d_S$) and overall rate of divergence ($d_N/d_S$) in *P. reticulata* based on Ensembl transcripts.** The X chromosome category excludes genes on the PAR and genes in Stratum I. 95% confidence intervals are based on bootstrapping with 1,000 replicates.

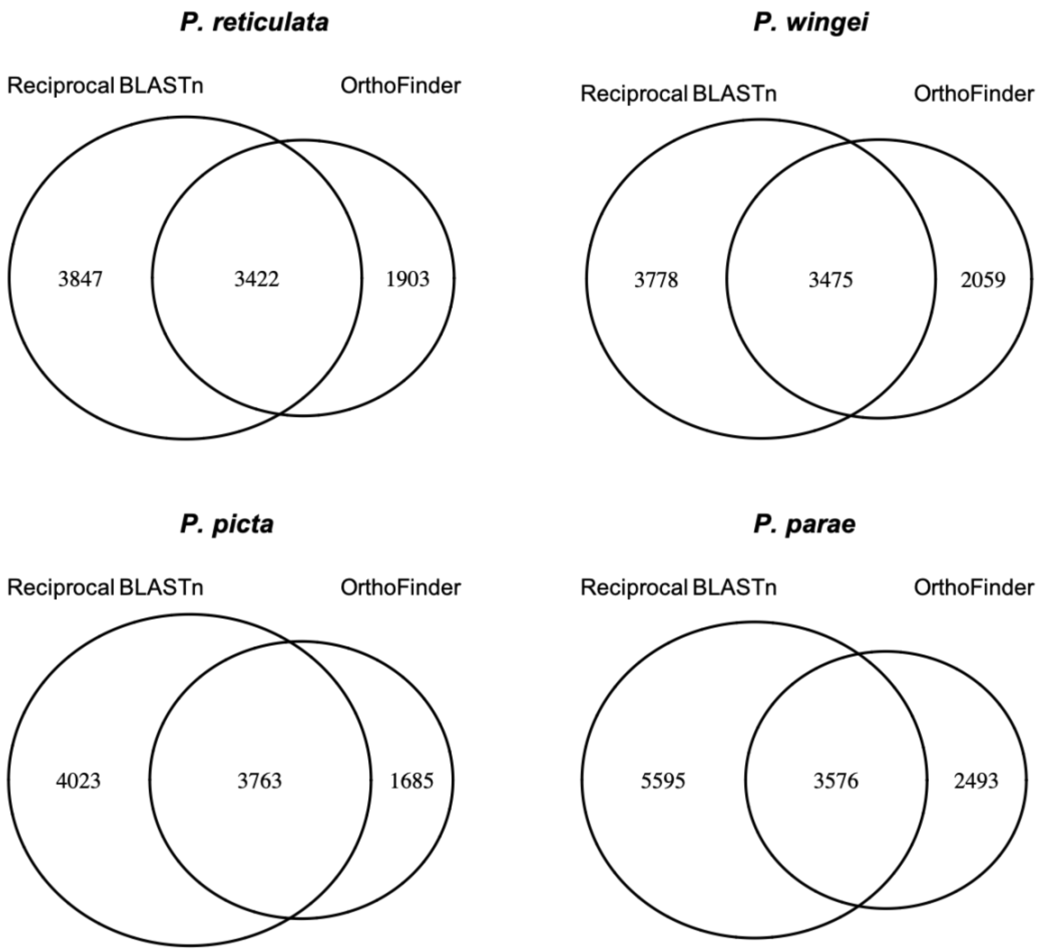


**Fig. S4. Number of orthogroups (four-way 1:1 orthologs) identified for the *P. reticulata*, *P. wingei*, *P. picta* and *P. parae* analyses using the reciprocal BLASTn and OrthoFinder approaches.**

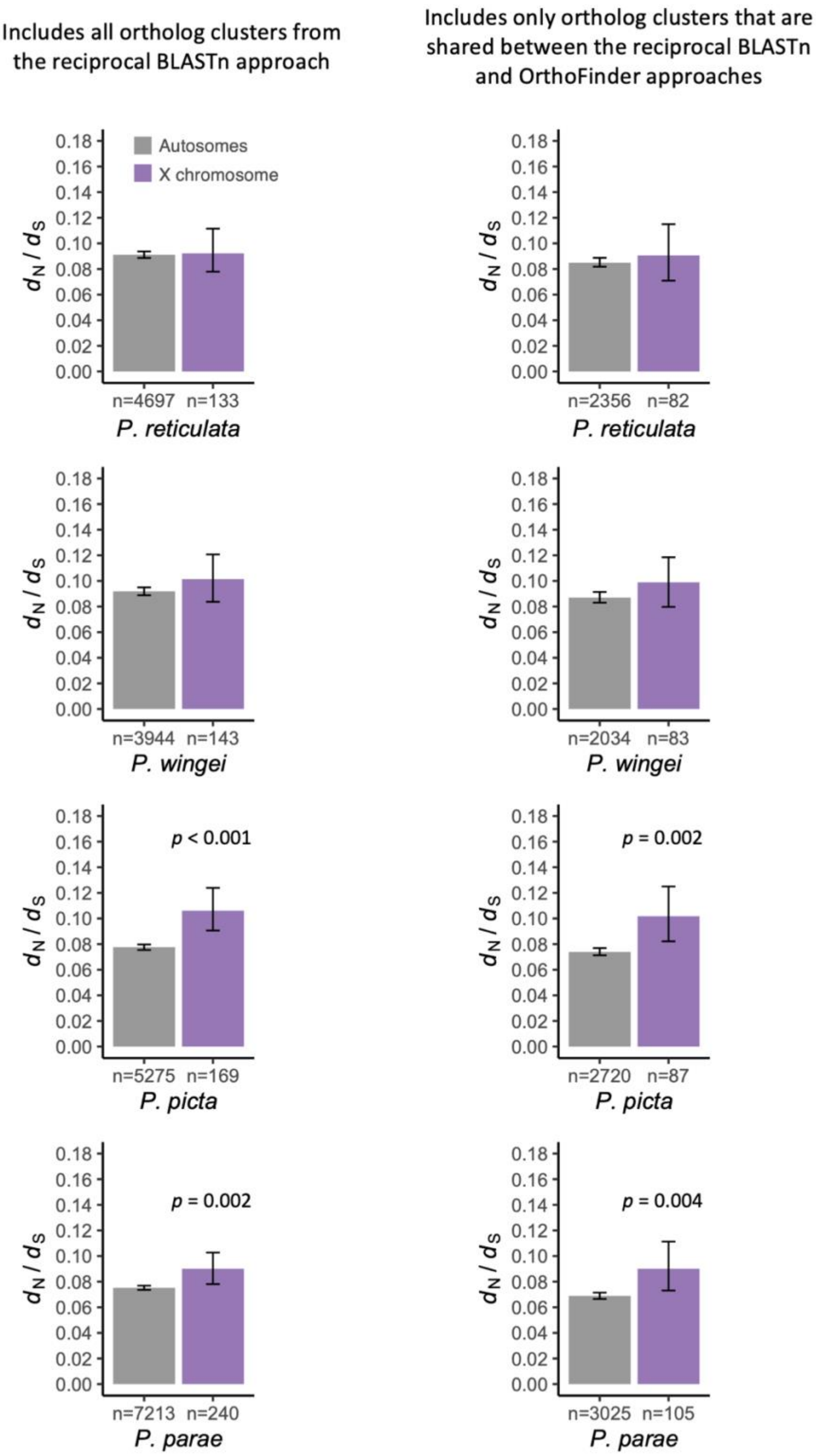


**Fig. S5. Estimates of divergence ($d_N/d_S$) for autosomal and X-linked genes in *P. reticulata*, *P. wingei*, *P. picta* and *P. parae* based on ortholog clusters from the reciprocal BLASTn approach (left) and based on ortholog clusters that are shared between the reciprocal BLASTn and OrthoFinder approaches (right).** In all cases the X chromosome category excludes genes on the PAR. 95% confidence intervals are based on bootstrapping with 1,000 replicates. Differences between autosomal and X-linked loci are based on 1,000 replicates permutation tests. Only significant differences are shown (*p* value < 0.01).
